# Restoring ancestral microbiome aids beetle adaptation to new diets

**DOI:** 10.1101/352989

**Authors:** Aparna Agarwal, Deepa Agashe

**Affiliations:** National Centre for Biological Sciences, Tata Institute of Fundamental Research, Bangalore, India

**Keywords:** Niche shift, Experimental evolution, Diet shift, Microbiome, Host-microbe association, Generalist

## Abstract

Eukaryotic hosts often depend on microbes that enhance their fitness, and such relationships may be relatively easily maintained in a stable environment. What is the fate of these associations under rapid environmental change? For instance, if the host switches to a new diet and/or encounters a different microbial community, how does the host-microbiome relationship change? Are the changes adaptive, and how rapidly do they occur? We addressed these questions with the red flour beetle *Tribolium castaneum*, a generalist insect pest that both consumes and lives in stored grain flour. We found that beetle fitness is enhanced by flour-acquired microbes in the ancestral habitat (wheat flour), but not in novel suboptimal environments (e.g. corn flour) that have a different resident microbial community. Beetles that disperse to new habitats thus have low fitness and a dramatically altered microbiome. Enriching novel habitats with ancestral (wheat-derived) microbes increased beetle fitness, suggesting a viable adaptive strategy. Indeed, within a few generations of laboratory adaptation to two distinct novel habitats, we found that beetle populations gradually restored their ancestral microbiome. Importantly, evolved populations showed a microbe-dependent increase in fecundity and survival on the new diet. We suggest that such repeated, rapid restoration of host-microbe associations may allow generalists to successfully colonize new habitats and escape extinction despite sudden environmental changes.

## INTRODUCTION

Host associated microbes can be crucial for host survival [1], as exemplified by a large body of work on insect-host associations. In particular, gut-associated microbial symbionts can provide their insect hosts with limiting nutrients [2] or enhance digestion of complex compounds [3]. For instance, bacteria in the midgut of the mosquito *Aedes aegypti* help lyse red blood cells, allowing efficient nutrient absorption by the host [4]. Microbes may also aid in detoxification of the host diet, as observed in bean bugs and stink bugs, whose symbionts degrade insecticides [3, 5]. Gut bacteria can thus directly influence host fitness: administering antibiotics reduces fecundity in *A. aegypti* [4] and delays larval growth in *Anopheles stephensi* [6]; and germ-free *Drosophila* show reduced lifespan and larval growth [7].

Such strong host-gut microbe associations are more likely to evolve in a stable environment [8], especially if microbes are transmitted across host generations. However, it is unclear whether and to what extent such insect-gut microbe interactions can be maintained when hosts experience significant environmental change within their lifetime, or across successive generations. For instance, generalist insects may feed on multiple resources in a few generations, potentially sampling a large diversity of diet-associated microbial communities. More generally, when any insect undergoes a dietary shift, both the host and its microbiome may face novel selection pressures. If the host benefits from its gut microbiome in the ancestral habitat, a dietary shift could disrupt the beneficial microbial community and reduce host fitness. In this scenario, what are the possible adaptive trajectories available to the host, and what is the effect on its microbiome?

Broadly speaking, there are four possible impacts of a host diet shift on the microbiome (Fig 1). For each case, there are two further possibilities: the host and its microbial community may be functionally associated, or the host may only passively acquire and house the microbial community. The likelihood of each of these trajectories depends on multiple factors, such as the difference between the nutritional content and fitness consequences of the ancestral and novel diets; the difference between the diet-associated microbes in each environment; and the relative rates of adaptive mutations in the host and microbes. However, to date no studies have directly tested the impact of environmental changes on the long-term fate of host-microbiome associations. As a specific example of rapid environmental change, dietary shifts present an opportunity to address this gap. Since dietary changes are ubiquitous across insects, examining host-microbiome interactions in the context of diet shifts may also offer new insights into insect ecology and evolution.

**Figure 1:**
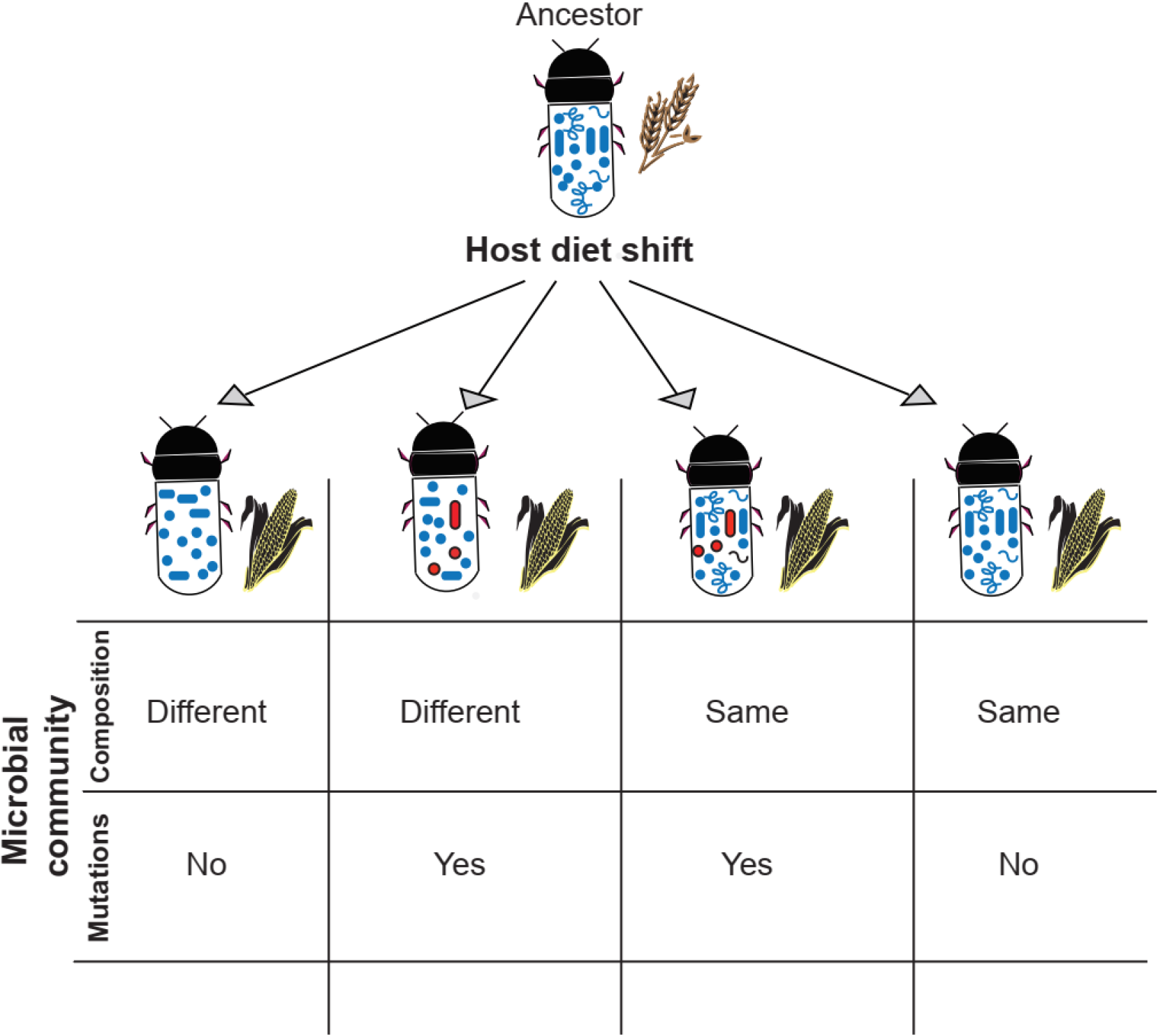
Possible impacts of a host diet shift on the composition and genetic makeup of the host-associated microbial community. Note that although microbes will most certainly acquire mutations during the course of the host diet shit, here we only refer to mutations that affect host fitness.

We addressed the role of microbes in novel habitats using the flour beetle *Tribolium castaneum*, a generalist insect pest that feeds on several cereal grain flours but is best adapted to wheat flour. All life stages of *T. castaneum* consume flour and excrete their wastes in the same habitat. Thus, gut microbes can easily spread and establish within a population, and a change in the dietary resource also represents a change in the environment and associated microbes. For our experiments, we used wild-collected beetles from stored wheat to establish an outbred population that exhibits maximum fitness on wheat flour. Thus, we refer to wheat as the ancestral resource, and we used corn, finger millet and sorghum – suboptimal diets with different nutritional content (Table S1) – as novel resources. 16S rRNA amplicon sequencing showed that each resource harboured distinct bacterial communities. We measured host fitness on each resource, either in the presence of normal flour-associated microbes or after depleting microbes with UV irradiation. Finally, we experimentally evolved beetle populations in two novel resources (corn and sorghum), and tested whether their fitness depended on new or ancestral flour microbes. We demonstrate rapid, adaptive restoration of host association with ancestral (wheat-derived) microbes, paving the way to understand the mechanistic basis of such generalist host-microbe associations.

## METHODS

### Beetle populations

For all experiments, we used an outbred population of the beetle *Tribolium castaneum*, generated using adults from 12 wild-collected populations from across India. We maintained stock populations in 4-week discrete generation cycles in wheat flour procured from a single company. We kept flour bags at −80 ° C for 4 h to kill any insect eggs in the flour. After this, we allowed the bags to thaw at room temperature. We used this flour for all our experiments. Each generation, we allowed adults to oviposit for 1 week and then removed them from the flour. After 4 weeks of development, we used resulting adult offspring to start the next generation. We housed populations in round plastic boxes with 2500 to 3000 adults per generation.

For experimentally evolving populations adapting to novel resources (13 populations in corn and 12 in sorghum), we founded replicate populations with 100, 200 or 500 adults at a density of 1g flour per individual (the populations were part of an independent study that required variable founding population sizes). These populations were also maintained in discrete generation cycles. Since larval development in corn is slow, we allowed corn populations 6 weeks for development, resulting in fewer generations of experimental evolution in corn. Each generation, we censused the number of live adult offspring and estimated per capita population growth rate as: (#Adults _t_ – #Adults _t−1_) / (#Adults _t−1_).

### Disrupting flour-associated microbial communities

We disrupted the flour microbial community by irradiating thin layers of flour under UV light for 2 h in a laminar hood without airflow. Alternatively, we mixed flour with one of three different broad spectrum antibiotics (tetracycline, ampicillin or kanamycin, 0.005% w/w). We added single eggs or adults to treated flour in a laminar hood (in 96-well microplates, Petridishes, microcentrifuge tubes, or boxes) and stored all containers in larger airtight boxes for the duration of experiment to prevent subsequent contamination. We handled and stored control groups (with untreated flour) under identical conditions, omitting only the irradiation step.

### Fitness assays

To successfully colonize a new environment, female fecundity and offspring survival are both critical. Therefore, to estimate fitness, we measured fecundity and egg survival in microbe depleted vs. untreated flour. To measure fecundity, we isolated 2 week old adult females in 0.7 g of sifted flour for 48 h, and counted the number of eggs laid by each female (n = 25 females per treatment). To measure egg survival, we collected ~100 individuals from the stock population, and allowed them to oviposit for 24 h in 100 g sifted wheat flour (sifting with a #50 sieve removes large flour particles, making it easier to identify and count eggs). We isolated eggs in 96 well plates and provided them with flour as required for each experimental treatment (e.g. untreated vs. UV-treated flour). We counted the number of surviving offspring after 3 weeks (n = 96 eggs/treatment/block; two independent blocks per treatment).

For experimentally evolved populations, we collected 50 females after the scheduled 1-week oviposition period of the 17th generation (for sorghum adapted populations) or the 10^th^ generation (for the corn adapted population). Recall that these females would have otherwise been discarded; hence, we did not disturb the evolving populations during these assays. The females were 2-3 weeks old at this stage, well within their peak fertility period. To measure fecundity, we isolated females and allowed them to oviposit in ~0.7 g sifted flour (untreated or UV treated corn or sorghum; n = 25 females/treatment) for 48 h. To measure egg survival, we collected ~100 individuals at the 18^th^ generation (sorghum adapted) or 11^th^ generation (corn adapted), and allowed them to oviposit in 50 g sifted flour for 24 h. We measured egg survival in the appropriate resource as described above. Evolved lines that did not successfully adapt to new resources had very low population size, and hence we did not have sufficient sample size to conduct fitness assays.

### Introducing microbes to UV treated flour

To analyse the fitness impact of microbes associated with ancestral or evolved populations, we introduced these microbes to UV-treated sifted flour via larval fecal matter. To test the impact of ancestral microbes, we allowed ~100 larvae (~two weeks of age) from a wheat stock population to consume and defecate in UV-treated flour for 24 h. To control for microbe-independent effects of introducing the larvae, we again treated half of this flour with UV to deplete the microbial load. We then measured fecundity in either microbe-enriched or depleted treatments, as described above (n = 25 females/treatment). To test the impact of microbes from adapted populations, we collected adults from the sorghum adapted population at generation 8, and allowed them to oviposit in fresh sorghum flour. We collected larvae after 2 weeks and used them to introduce sorghum-associated microbes in UV-treated wheat or sorghum. Similarly, we collected adults from the corn adapted population at generation 12, allowed them to oviposit in fresh corn flour, and used 2 week old larvae to introduce corn adapted microbes in UV treated corn.

### Determining bacterial community composition

We determined the bacterial community associated with flour samples and beetles using amplicon sequencing of the 16S rRNA gene. For each treatment, we isolated 2-7 individuals (larvae or 1 week old adult females) and surface sterilized them using 70% ethanol, followed by a wash with DNAse and RNAse-free ultra-pure water. To identify flour-associated microbes, we collected four replicate samples of flour (~0.07 g each). We extracted DNA from each sample using the Promega DNA extraction kit, following the manufacturer’s instructions for extracting bacterial DNA. To minimize protein contamination from beetle tissue or flour, we increased incubation time with proteinase K from 3 h to 12 h. We prepared barcoded 16S libraries by PCR amplification with standard 16S Illumina primers, and KAPA HiFi Hotstart mix. We sequenced libraries on the Illumina Miseq platform (300 bp paired end sequencing), following the Illumina protocol for further amplification and cleanup steps. We used standard QIIME pipelines [9–13] to generate tables with the relative abundance of all OTUs (Operational Taxonomic Units with 97% sequence similarity), assigning taxonomy using closed reference OTU picking with the Greengenes database [14]. We removed chloroplast or mitochondrial reads using the filter_taxa_from_OTU_table.py command in QIIME (Figs S1-S3). To avoid rare OTUs that may represent contamination, we removed OTUs represented by less than 20 reads. We did not get any detectable amplification in a negative control sample (ultra-pure water through the DNA extraction protocol) as measured by Qubit HS assay, at all the PCR steps, suggesting that indeed contamination levels in our sequencing method are low. After this filtering, we found that some samples did not have any bacterial OTUs, and we removed these from further analysis (Table S2). Note, however, that these are also informative samples and we discuss them while presenting our results. We used the final set of samples and OTUs to re-calculate the relative abundance of each OTU per sample. We carried out all subsequent analysis in R version 3.2.2 [15].

On average, we found ~300 bacterial OTUs in flour samples and ~450 bacterial OTUs in beetle samples (Figs S1-S3). We first visualised the entire bacterial community present in each sample using an unconstrained clustering approach (Principle Co-ordinate analysis, PcoA) with the pcoa function in the R package ape v5.1 [16], and generated plots using the biplots function in the R package BiplotGUI [17]. We also used constrained clustering (ordination analysis) with the CAPdiscrim function in the BiodiversityR package [18]. To statistically test the impact of flour and resource treatments on full bacterial communities, we used PERMANOVA analyses, implemented with Adonis function in the package Vegan [19].

The full bacterial community is complex and is hence difficult to visualise. Hence, to visualize variation in the most dominant bacterial OTUs, we also analysed the five most abundant OTUs across replicate samples of a given treatment (see Fig S4 for an example). These dominant bacteria are more likely to play a functional role in the host-microbe association, since we expect beneficial bacteria to be enriched in the beetle-associated community. Note that various samples within a comparison set may not share any of their most abundant OTUs. Hence, in the final list of most abundant OTUs in a specific comparison, we could have anywhere between 5 OTUs (if all abundant OTUs were shared) and 25 OTUs (if the five most abundant OTUs were unique for each group). We clubbed all bacterial OTUs that were not amongst the 5 most abundant in any sample into the category “others”.

## RESULTS AND DISCUSSION

Darwin famously started his landmark paper to the Linnaean society [20] with the phrase “All nature is at war”, referring to organisms’ continued struggle for existence. However, many eukaryotes are not alone in this struggle, but rely on microbes that provide crucial fitness benefits. However, most of our understanding of such host-microbe relationships derives from studies in single environments. What happens when organisms disperse to new habitats, their microbiome is disrupted, and their microbial partners are either missing or cannot establish in the new environment?

### Beetle fitness depends on flour microbes in the ancestral, but not novel resource

We first tested whether flour beetles derive a fitness benefit from their microbial flora, and whether the microbes are environmentally acquired or vertically transmitted. We disrupted the microbial community associated with the flour beetles’ environment by treating flour with UV radiation, and introduced individuals to treated flour. Within 48 h, we observed a significant reduction in female fecundity in UV-treated wheat flour (Fig 2A; t-test for UV-treated vs. untreated wheat flour, p =0.0002. Similarly, across much longer timescales (3 weeks), egg survival also decreased in UV-treated wheat (Fig 2B; ChiSq test for count data, p = 0.003), as did other fitness proxies such as adult lifespan and body mass (Fig S5; Kaplan-Meier test for lifespan: p <0.01; t-test for body mass, p <0.01). We observed a similar reduction in beetle fitness when we mixed broad-spectrum antibiotics in wheat flour (Fig S6A-B; ANOVA for the effect of antibiotics: p <0.01, ChiSq test: p <0.01, df =7). These results suggest that flour beetle fitness depends on flour-acquired microbes that are not maternally transmitted. In contrast to the patterns in wheat, we found that UV treatment had no impact on beetle fitness in three novel resources (corn, sorghum and finger millet; t-test for each flour, p > 0.05; Fig 2A-B), and adding Ampicillin to Sorghum flour did not alter fecundity in Sorghum (t-test for the effect of ampicillin in sorghum, p = 0.99; Fig S6C). Thus, beetle fitness depends strongly on flour-associated environment microbes at both larval and adult life stages, but only in the ancestral wheat resource to which the hosts are well adapted.

**Figure 2:**
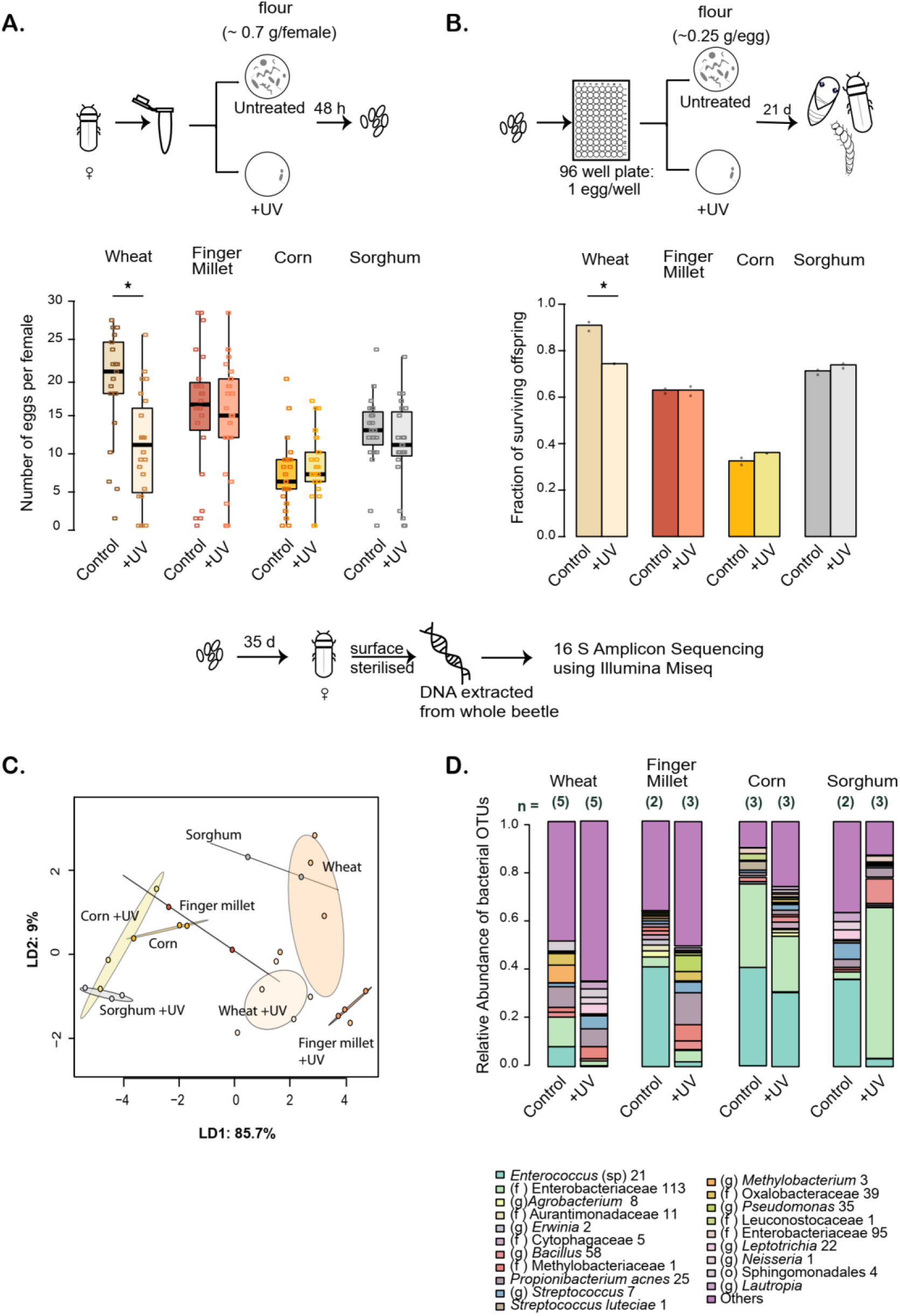
Flour beetle fitness depends on flour-associated microbes in the ancestral resource. Each panel includes a schematic representation of relevant experiments. **(A)** Fecundity (number of eggs laid per female; n = 25 females per treatment) in untreated or UV-treated ancestral (wheat) or novel resources (finger millet, corn and sorghum). Boxplots show the median number of eggs laid in each treatment (boxes indicate inter quartile length (IQL) and whiskers indicate 1.5× IQL). Raw data points are shown as open squares. Asterisks indicate a significant difference between untreated and UV-treated resource. **(B)** Average fraction of surviving offspring after 3 weeks of development in each resource (n = 96 stock-collected eggs per treatment per block; 2 blocks per treatment; open circles show the fraction of survival/treatment for each independent block. Asterisks indicate a significant difference between untreated and UV-treated resource. **(C)** Linear discriminant (LD) analysis of the complete bacterial communities associated with individuals reared on different resources. LD1 and LD2 are the first two discriminants that best capture the classification of the different groups; percent variation explained is given in parentheses. Each filled circle represents an individual beetle, and ellipses indicate 95% confidence intervals. **(D)** Dominant bacterial community members associated with beetles fed on different flours. Stacked bar plots show the average relative abundance of the 5 most abundant bacterial OTUs from individual beetles (sample size is given above each bar). OTUs were classified to the lowest taxonomic level possible, indicated in parentheses (o=order, f=family, g=genus). Numbers after OTU names distinguish OTUs with the same taxonomic classification that were distinct at sequence level (97% identity).

Why is the fitness impact of flour microbes resource-dependent? One possibility is that the novel resources are so suboptimal that we could not detect a small impact of flour microbes in these assays. However, this is unlikely because beetle fitness in finger millet and sorghum is not dramatically different from that in untreated wheat (Fig 2A-B). Thus, our results probably do not reflect the strength of selection imposed by the novel environment. Another possibility is that we did not disrupt the microbial community in the novel resources sufficiently, and therefore we did not observe a fitness effect. However, 16S rRNA amplicon sequencing showed that different flours harbour distinct bacterial communities, and that UV treatment significantly disrupted the communities in each case (Fig S7; PERMANOVA: p_resource_ <0.01; p_UV_ <0.01; p_resourcexUV_<0.01). Importantly, we observed similar patterns for microbiomes of beetles that fed on these resources (PERMANOVA: p_resource_ = 0.001; p_UV_ = 0.314; p_resourcexUV_ <0.01). Individuals consuming untreated vs. UV treated flours formed distinct clusters in a linear discriminant plot (Fig 2C; see Fig S8 for unconstrained PCoA), despite substantial variation across host individuals and the lack of detectable bacterial reads in a few beetles (Table S2). The difference in microbiomes of beetles reared on different flours, and the impact of UV treatment, was especially striking when we focused on dominant bacterial OTUs across treatments (Fig 2D). Finally, we observed that the microbiome of wheat-reared beetles is distinct from the microbiome of wheat flour (Fig S7; PERMANOVA: p_sample type_ = 0.001), indicating that only specific bacteria colonize the beetle gut, and the entire flour-associated community is not passively harboured. Thus, our results show that beetle fitness depends on beneficial microbes found in wheat, and that resource-specific fitness impacts of microbes arise because the novel flours are associated with a distinct set of microbes.

### Ancestral microbiome is also beneficial in novel environments

Since beetles depend on wheat-derived microbes, we hypothesized that these “ancestral” microbes may also provide a fitness benefit in the novel habitats. Thus, we predicted that enriching the novel environment with wheat-associated microbes should improve beetle fitness in novel resources. To test this, we focused on corn and sorghum, which imposed low fitness relative to wheat. We briefly introduced wheat-fed larvae into each novel resource, so that the larvae would add their fecal matter to the flour and enrich it with wheat-associated microbiomes. We likely introduced very high bacterial loads through larvae because we used a large number of larvae, whose guts turn over a very high volume of food. Enriching corn and sorghum with wheat microbes caused a significant increase in fecundity (Fig 3). Importantly, beetle fecundity decreased in enriched flour treated with UV, confirming that the observed impact on fitness is due to microbial enrichment rather than other larval secretions (t-test: p_UV_ <0.01). Ancestral microbes also rescued egg survival in corn, although they could not rescue survival in sorghum (Fig S10). Thus, ancestral microbes could provide a fitness advantage even in novel environments, and maintenance of the ancestral microbiome could be a viable adaptive strategy after dispersal to new habitats. However, as described above the adult beetle microbiome is dramatically altered immediately after introduction to new resources (Fig 2), and it is not clear whether this challenge could be overcome during the course of adaptation to new habitats.

**Figure 3.**
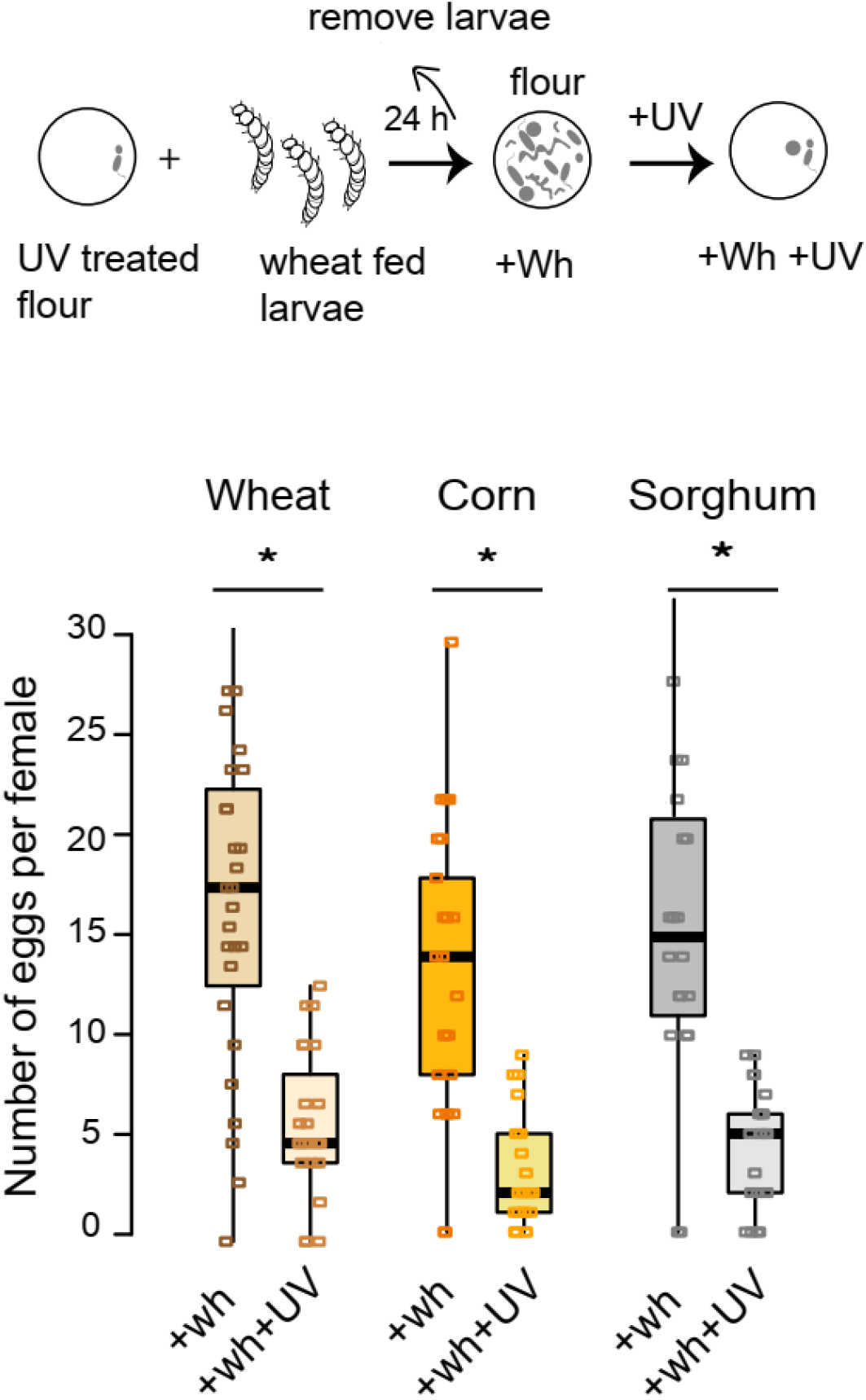
Introducing ancestral microbes in novel resources increases fecundity. Boxplots show the median number of eggs laid in flour enriched with wheat microbes, or in enriched flour after further UV treatment (see schematic on top; +Wh = wheat microbes; n = 25 females per treatment; boxplots and raw data are shown as described in Fig 2). Asterisks indicate a significant difference between untreated and UV-treated resource.

### Gradual restoration of ancestral microbiome aids adaptation to new resources

We hypothesised that co-habitation between wheat microbes and beetles for several generations may have facilitated beetle dependence on wheat microbes. We therefore predicted that as beetle populations adapt to novel environments, there may gradually enrich the ancestral microbial community, with or without beneficial microbial mutations (Fig 1). Alternatively, beetles could establish a novel relationship with corn- or sorghum-specific microbes; or adapt to the new resources independently of flour microbes. Fortunately, for an independent project, we had previously allowed replicate beetle populations to evolve under selection in corn (13 populations) or sorghum (12 populations) (Fig 4A). The populations were founded with wheat-reared adult beetles, who may have introduced wheat-associated microbes in the novel habitat through their fecal matter. Hence, we used these populations to distinguish between the possibilities outlined in Fig 1. Within 10-15 generations, four populations showed a clear positive growth rate (one in corn and three in sorghum; Fig 4B-C). We tested whether beetles from these “adapted” populations showed an association with flour microbes, and whether the association was beneficial. Where possible (see methods), we also analysed one population from each resource that had avoided extinction, but did not have a positive growth rate (“not adapted” populations; Fig 4B-C).

**Figure 4.**
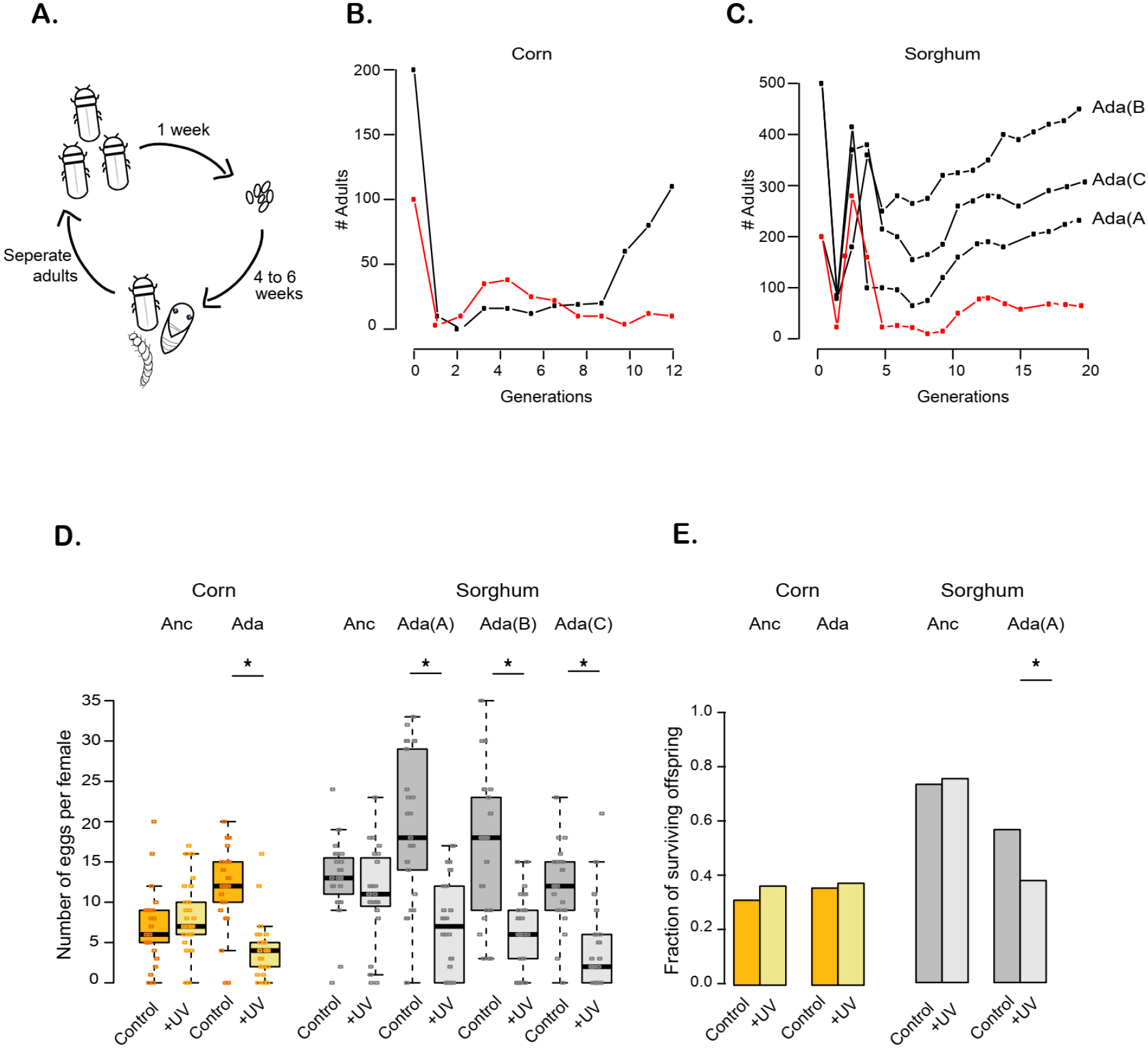
Fitness of beetles adapted to novel resources depends on flour-associated microbes. **(A)**A schematic representation of the experimental evolution regime. **(B and C)** Number of adults per population across generations, for adapted populations (black) and one population that did not successfully adapt (red), for **(B)** corn and **(C)** sorghum. **(D)** Fecundity (n = 25 females per treatment) in untreated vs. UV-treated flour for ancestral females (“Anc”: fed on the novel resource for a single generation), or for females from populations that had adapted to the novel resource (“Ada”). Boxplots and raw data are shown as described in Fig 2. Asterisks indicate a significant difference between untreated and UV-treated resource. **(E)** Fraction of surviving offspring after 3 weeks of development in untreated or UV-treated flour (n = 96 stock-collected eggs per treatment), for eggs derived from ancestral or adapted populations as above. Asterisks indicate a significant difference between untreated and UV-treated resource.

We found multiple lines of evidence suggesting that gradual restoration of ancestral (wheat-derived) microbiomes aided adaptation to novel resources. Whereas beetle fitness was unaffected by flour microbes immediately after introduction to new resources, all adapted populations derived a fitness advantage from environmental microbes (Fig 4 D-E): beetle fecundity was lower in UV treated flour (Fig 4D; pairwise t-test for each adapted population for the effect of UV treatment: p < 0.01), and in one of the adapted sorghum populations, egg survival was also microbe-dependent (Fig 4E; ChiSq test for the effect of UV treatment. In corn, p>0.05; In sorghum p_Anc_ = 0.8, p_Ada(A)_< 0.01). Importantly, microbiomes of beetles from adapted populations were similar to that of wheat-reared ancestors, but distinct from the bacterial community of individuals fed on sorghum or corn for a single generation (Fig 5A-B; PERMANOVA p_(Ancestor(novel resource) vs. Adapted(novel resource)_ = 0.003; p_(Ancestor(wheat) vs. Adapted(novel resource)_ = 0.31; see Fig S11 for PCoA). This pattern of congruence in the microbiome is especially clear if we focus on dominant bacterial taxa (Fig 5C). Conversely, in populations that did not adapt successfully, beetles harboured bacterial communities that were similar to ancestral individuals fed on the respective flour for a single generation (Fig 5A-C; PERMANOVA p_(NotAdapted(novel resource) vs. Ancestor(novel resource)_ > 0.1 for both corn and sorghum). Finally, we found that the microbiomes of adapted populations were functionally similar to the ancestral microbiome, such that microbes from a sorghum-adapted population could rescue ancestral beetle fitness in wheat flour (Fig 5D). Microbes from sorghum or corn-adapted lines also elicited a microbe-dependent fecundity response in ancestral females exposed to the respective resource (Fig 5D, t test for the effect of resterilizing the flour with UV: p_wheat_<0.01; p_sorghum_ = 0.006; p_Corn_ = 0.002), mimicking the effect of ancestral wheat microbiomes (although the magnitude of the effect was lower; compare Fig 5D with Fig 2). Thus, despite new mutations that probably occurred in hosts and bacteria during experimental evolution, the “evolved” host-bacterial association was effectively equivalent to the ancestral microbiome. We note that although all aspects of host fitness are not explained by the host-microbial association, it is clear that restoring ancestral partnerships with bacteria played an important role during adaptation in four independently evolved beetle populations.

**Figure 5:**
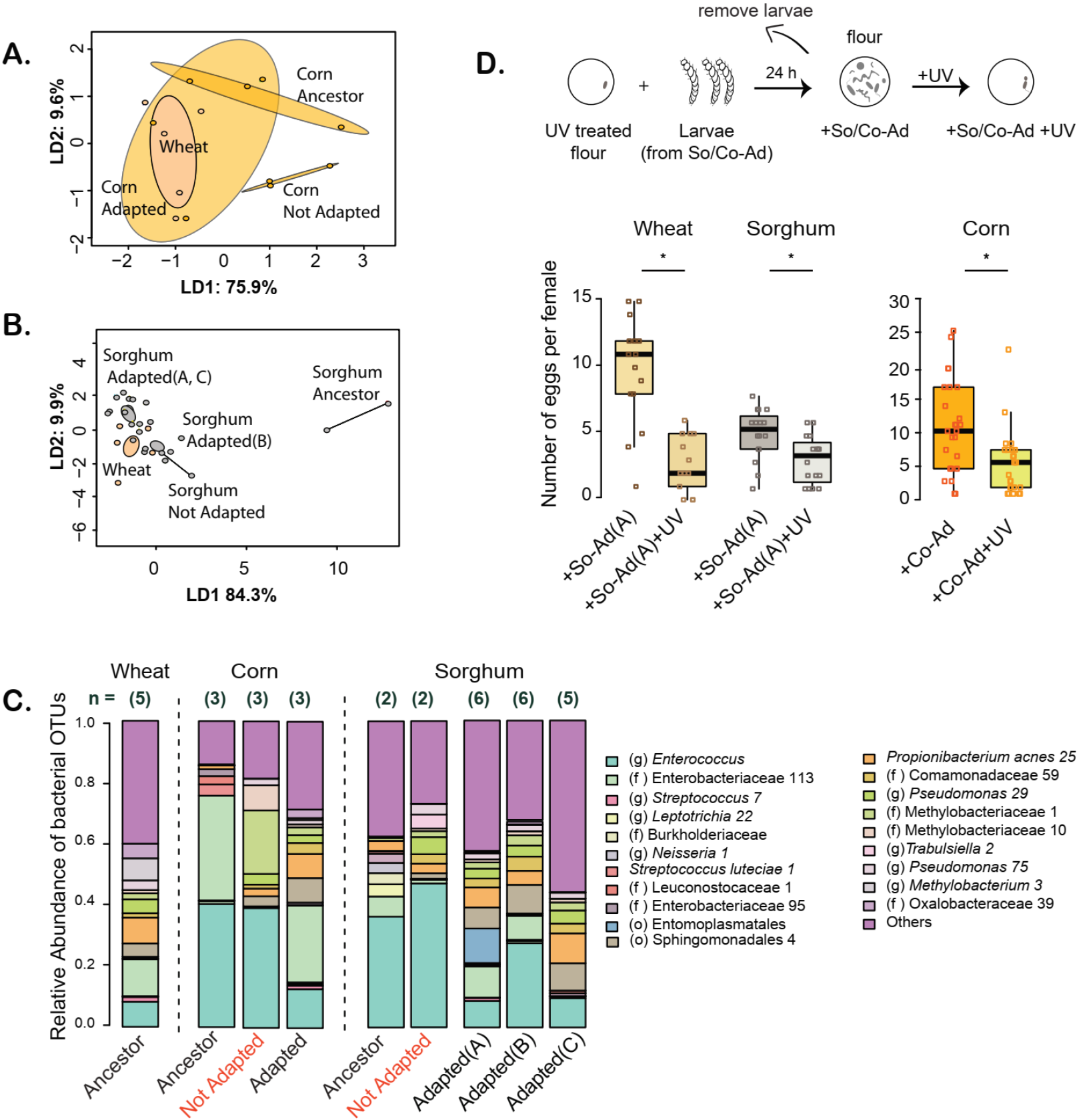
Beetles from adapted populations converge on the ancestral wheat-associated microbiome. **(A, B)** Linear discriminant analysis for full bacterial communities associated with individuals from adapted vs. not adapted populations, including wheat-fed ancestral individuals (panel A: corn; B: sorghum). Discriminants LD1 and LD2 best capture the classification of different groups, as indicated by the percent variation explained by each. Filled circles indicate individual beetles and ellipses show 95% confidence intervals. **(C)** Dominant members of the bacterial community of beetles from adapted vs. not adapted populations, and their respective ancestors. Stacked bar plots show the average relative abundance of the five most abundant bacterial OTUs in individual beetles (sample size indicated above bars) fed on untreated or UV treated flour. OTUs were classified to the lowest taxonomic level possible, indicated in parentheses (o=order, f=family, g=genus). Numbers after OTU names distinguish OTUs with the same taxonomic classification. **(D)** The impact of microbes from corn (Co-Ad) or sorghum-adapted (So-Ad) populations on the fecundity of wheat-adapted females (n = 25 females per treatment; see schematic on top). The left panel shows fecundity after 24 h in sorghum or wheat flour enriched with microbes from a sorghum adapted population (So-Ad), vs. enriched flour that was again treated with UV (So-Ad+UV). The right panel shows fecundity measured after 48 h in corn flour enriched with microbes from the corn adapted population (Co-Ad), vs. in enriched flour treated with UV (Co-Ad+UV). Boxplots and raw data are shown as described in Fig 2. Asterisks indicate a significant difference between untreated and UV-treated resource.

### A novel yet simple adaptive path in new habitats

Based on our results (summarized in Fig 6), we postulate the following trajectory of changes in host-microbiome association during adaptation. Immediately after introduction to the novel habitats, the microbiome of founding beetles shifted dramatically to reflect the bacterial community associated with the new diet. Although the founding adults also carried (and introduced) wheat-derived bacteria via their fecal matter, these bacteria were either rare or were unable to effectively colonize the beetle gut in the presence of the novel flour-associated microbes and the new diet. Thus, initially very few beetles harboured beneficial bacteria. Over generations – because individuals that harboured the bacteria also reproduced more – the abundance of the ancestral beneficial bacteria increased in the new habitat. At this stage, we could observe a dependence of beetle fitness on the flour microbes.

**Figure 6:**
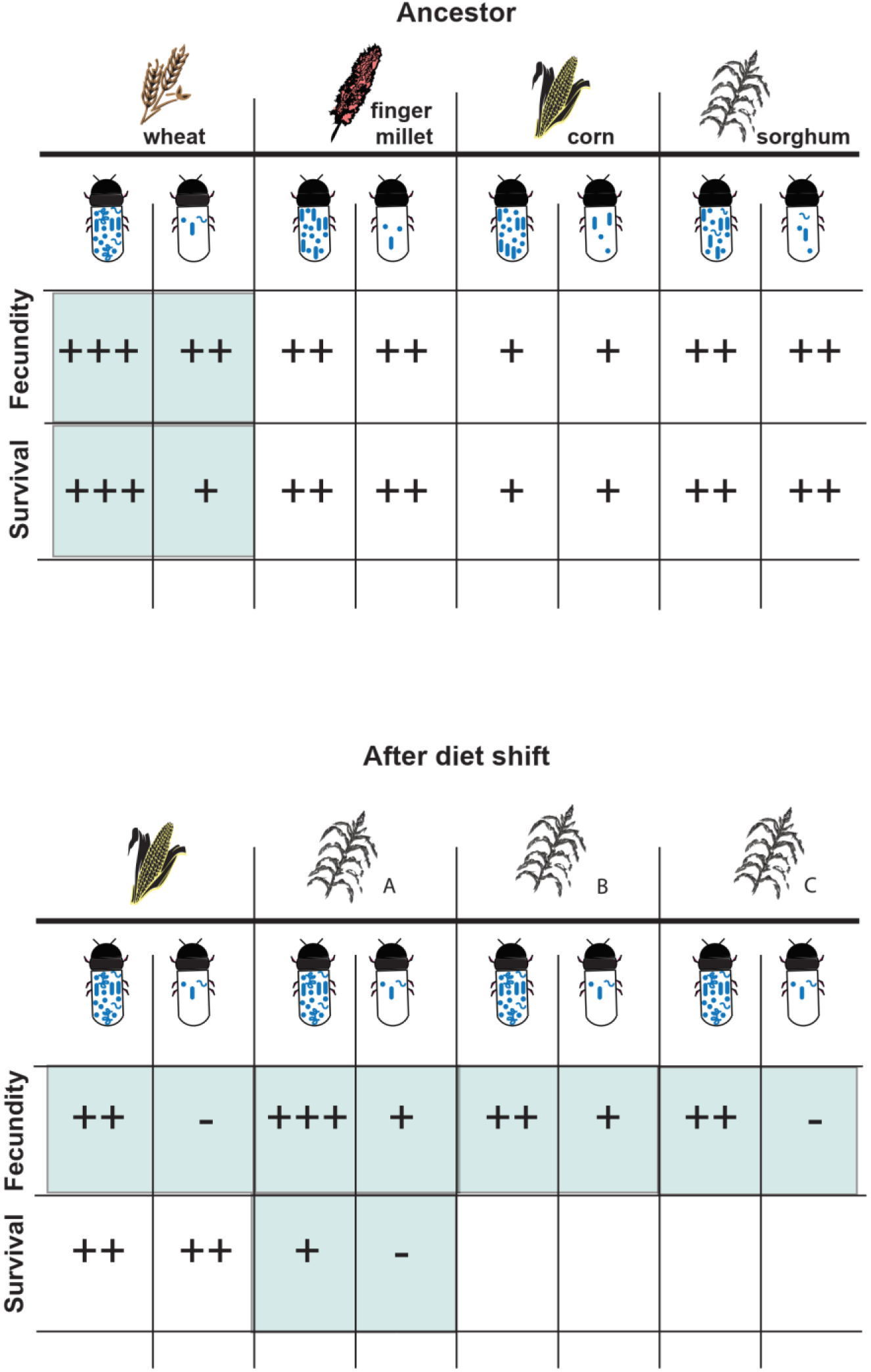
Summary of patterns of change in host dependence on microbes during experimental diet shifts. “+” signs indicate relative fitness of beetles in each resource, with or without access to flour-associated microbes (as indicated by bacteria inside beetles). Instances where we observed a significant microbe dependent fitness decline are highlighted in pale green (based on pairwise comparison between beetles with normal vs. depleted microbial loads). Empty cells indicate missing data.

Our results suggest that “ecological” changes in microbial community composition may be sufficient to facilitate host adaptation to novel habitats. Although the bacteria almost certainly acquired genomic mutations during experimental evolution, our results indicate that such genetic changes may not be critical in the early stages of host adaptation. Over longer evolutionary timescales, it is possible that bacteria would acquire flour- or host population- specific beneficial mutations, such that ancestral beetles would not benefit from the microbiomes of evolved beetles. Note that although the bacterial communities associated with the wheat ancestor and the adapted populations are structurally similar, the same bacterial taxa dominated communities of beetles from non-adapted populations as well as naïve beetles fed on the novel resources for a single generation. For instance, two major bacterial genera – *Enterococcus* and *Enterobacteriacae* – were associated with all sampled beetles (Fig 2), potentially reflecting a superior ability to colonize the beetle gut. However, mere colonization by these bacteria is not sufficient to provide fitness advantages, since beetle fitness was initially low in the novel environments. Instead, the relative and/or absolute abundances of other bacteria may also be important for host fitness. Further experiments to selectively add or eliminate specific bacteria, in combination with deep sequencing of the microbiome, are necessary to test this possibility.

Since beetles converged on similar communities in different resources, we also speculate that bacteria associated with such generalist beetles may themselves be generalists, enhancing host fitness across multiple resources. Indeed, bacteria from the genus *Enterococcus* and family Enterobacteriacae (dominant taxa associated with wheat–adapted beetles) are frequently found in the guts of several insects [8]. In general, Enterococci can utilize a variety of sugars and carbohydrates; for instance, the human gut commensal *Enterococcus faecalis* can digest a wide range of plant based carbohydrates such as cellulose, which humans cannot digest [21, 22]. Similarly, members of the family Enterobacteriacae are commonly found in stored grain warehouses, and can grow on a variety of cereal grains [23]. Thus, these bacteria may have the metabolic potential to utilise multiple resources. In further work, we hope to test whether these taxa are particularly suited to colonizing the beetle gut and/or use various cereal grains, and are specifically responsible for the observed beetle-microbiome association.

## Conclusions

We observed surprisingly rapid and repeatable restoration of ancestral microbiomes across different resources and populations, suggesting a fascinating paradigm for host evolution in new habitats. We propose that this may be a general phenomenon whereby introducing ancestral microbes can reduce the probability of host extinction in a novel environment. Conversely, host-mediated microbial dispersal may also allow bacteria to colonize diverse habitats, while significantly changing the microbial communities in new environments. Therefore, both the bacterial partners and the host may impact each other’s ability to sample and colonize new environments. Our study system thus presents a unique opportunity to analyse hosts as well as their associated bacteria during the establishment of host-microbial associations.

## ACKNOWLEDGEMENTS

We thank members of the Agashe lab for comments on the manuscript; Amruta Rajarajan, Soumya Panyam, Gaurav Agavekar and Riddhi Deshmukh for laboratory assistance; Gaurav Diwan, Kruttika Phalnikar, Umesh Mohan and Rittik Deb for help with QIIME and R analysis; and Awadhesh Pandit and other staff from the Next Generation Sequencing facility at NCBS. We acknowledge funding and support from the Wellcome Trust DBT India Alliance Fellowship (grant number IA/I/17/1/503091 to DA) and the National Centre for Biological Sciences (NCBS).

## AUTHOR CONTRIBUTIONS

Conceived and designed experiments: DA, AA. Conducted experiments: AA. Analysed data: AA, DA. Wrote the manuscript: DA, AA. Acquired funding: DA.

